# Keratinocyte Piezo1 drives paclitaxel-induced mechanical hypersensitivity

**DOI:** 10.1101/2023.12.12.571332

**Authors:** Alexander R Mikesell, Elena Isaeva, Marie L Schulte, Anthony D Menzel, Anvitha Sriram, Megan M Prahl, Seung Min Shin, Katelyn E Sadler, Hongwei Yu, Cheryl L Stucky

## Abstract

Recent work demonstrates that epidermal keratinocytes are critical for normal touch sensation. However, it is unknown if keratinocytes contribute to touch evoked pain and hypersensitivity following tissue injury. Here, we used inhibitory optogenetic and chemogenetic techniques to determine the extent to which keratinocyte activity contributes to the severe neuropathic pain that accompanies chemotherapeutic treatment. We found that keratinocyte inhibition largely alleviates paclitaxel-induced mechanical hypersensitivity. Furthermore, we found that paclitaxel exposure sensitizes mouse and human keratinocytes to mechanical stimulation through the keratinocyte mechanotransducer Piezo1. These findings demonstrate the contribution of non-neuronal cutaneous cells to neuropathic pain and pave the way for the development of new pain-relief strategies that target epidermal keratinocytes and Piezo1.

**Summary:** Sensitization of the keratinocyte mechanotransducer Piezo1 drives paclitaxel-induced touch pain.

## INTRODUCTION

Paclitaxel is a primary therapeutic intervention for breast, lung, and ovarian cancers^1^. Paclitaxel inhibits tumor cell proliferation and growth primarily through stabilization of microtubules. Despite its impressive response rate of up to 86%, its use is marred by side effects such as chemotherapy-induced peripheral neuropathy (CIPN)^2^. CIPN affects 60-70% of paclitaxel-treated patients and causes painful sensory symptoms such as severe hypersensitivity to skin touch and temperature, dysesthesia (pins and needles), and unprovoked burning or shooting-like pain. CIPN is often so severe that it necessitates dose reduction or cessation of treatment^3^. There is currently no Food and Drug Administration approved treatment for managing CIPN. Commonly, patients are prescribed opioids, antidepressants, and anti-seizure medicines, but these options have limited effectiveness and a range of adverse side effects. Thus, the development of novel therapeutics for CIPN is desperately needed.

Paclitaxel-induced pain has been attributed to accumulation of the drug in the dorsal root ganglia and spinal cord, where it damages peripheral and central neurons^3^. However, paclitaxel also accumulates at high levels in the skin, often leading to severe dermatological reactions, such as inflammatory rashes, blistering lesions, and hyperpigmentation^4–6^. Despite this known accumulation, the specific impact of paclitaxel on the initial transduction and/or amplification of sensory stimuli in the epidermis remains unexplored. The epidermis is the most superficial layer of skin that covers nearly the entire mammalian body. Over 95% of cells in the epidermis are keratinocytes. Recent evidence demonstrates that keratinocytes are critical for normal touch sensation^7–9^. Keratinocytes respond directly to mechanical stimulation through expression of Piezo1, a bona fide mechanically gated ion channel necessary for primary afferent and behavioral responses to tactile stimuli^10,11^. Upon mechanical stimulation, keratinocytes release adenosine triphosphate (ATP) that activates purinergic receptor P2X4 on neighboring sensory neurons, many of which form “synapse-like” connections with keratinocytes^9,12^. Skin injury in many forms leads to pain and hypersensitivity to both gentle and noxious tactile stimuli, but whether keratinocytes play a role in cutaneous pain after tissue or nerve injury is unknown. If keratinocytes contribute to driving CIPN-related pain, they may be targeted in the future by non-invasive pain treatments that alleviate pain superficially and avoid the central nervous system side effects of many current analgesics, potentially allowing patients to receive longer or more aggressive cancer treatments. Here, we tested the hypothesis that keratinocytes mediate paclitaxel-induced mechanical pain through the mechanotransducer Piezo1. Together these data are the first to show a direct role of keratinocytes in driving paclitaxel-induced mechanical hypersensitivity.

## RESULTS

### Keratinocyte inhibition alleviates paclitaxel-induced mechanical hypersensitivity

We began investigating whether keratinocyte activity contributes to paclitaxel-induced mechanical hypersensitivity by selectively inhibiting keratinocytes optogenetically. We used mice that exclusively express GFP-tagged Archaerhodopsin-3 (Arch) in Keratin-14 (K14) expressing epidermal cells (Arch-K14Cre+) (**Fig. S1A**). To optogenetically inhibit keratinocytes, we exposed the plantar surface of each hindpaw to amber light (590 nm) 1 min before and during von Frey stimulation. This brief optogenetic inhibition largely reversed the paclitaxel-induced mechanical hypersensitivity in Arch-K14Cre+ mice whereas mechanical responses of Arch-K14Cre-littermate controls were unaffected (**Fig. 1A**). Note that a similar extent of mechanical hypersensitivity reversal was observed with keratinocyte inhibition at all time points tested (2 days – 3 weeks post initial paclitaxel injection) (**Fig 1A**). Optogenetic keratinocyte inhibition also increased the withdrawal thresholds of naïve, uninjured Arch-K14Cre+ mice, similar to previous reports^9^. The effect of keratinocyte inhibition in paclitaxel-treated Arch-K14Cre+ mice was greater than that in uninjured Arch-K14Cre+ mice, suggesting that the contribution of keratinocyte activity to mechanical sensitivity *in vivo* is greater after paclitaxel injury than in healthy tissue (**Fig 1B**). The robust effect of keratinocyte inhibition was transient because mechanical responses returned to baseline withdrawal thresholds one minute following the removal of the 590 nm laser (**Fig. S1B**). This suggests that keratinocyte activity is required for paclitaxel-induced mechanical hypersensitivity. To determine whether keratinocytes also contribute to other forms of behavioral mechanical hypersensitivity besides innocuous punctate von Frey stimulation, we tested dynamic hypersensitivity using a delicate paint brush swipe and noxious pinprick at 10 days after the start of paclitaxel treatment. The heightened attending responses to both paintbrush and pinprick induced by paclitaxel were substantially reduced in the presence of brief inhibitory light in Arch-K14Cre+ mice, but not in Arch-K14Cre-littermate controls (**Fig. 1C, D**).

**Fig. 1.**
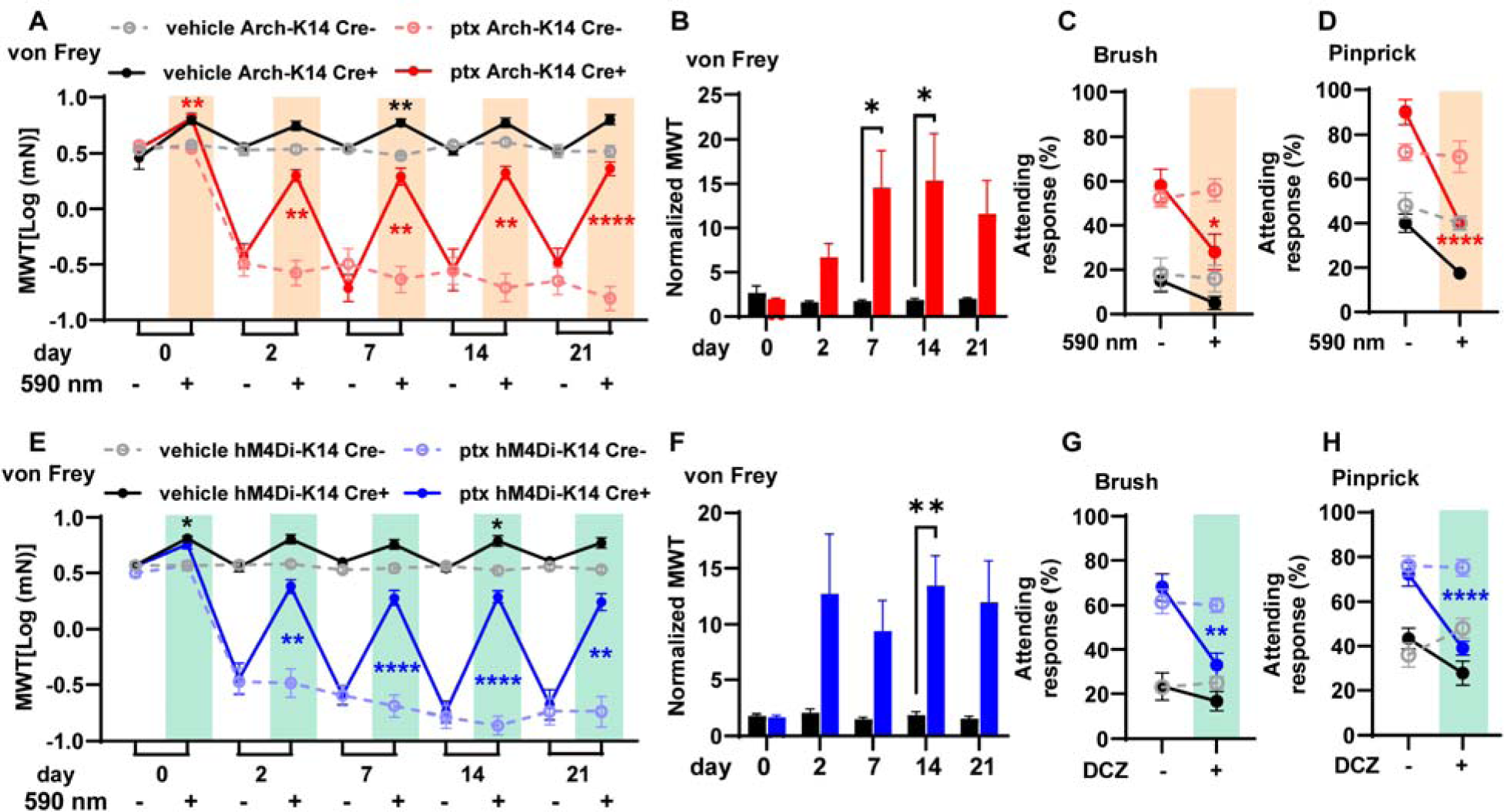
Alleviation of paclitaxel-induced mechanical hypersensitivity through optogenetic and chemogenetic inhibition of keratinocytes: (**A**) Comparison of mechanical withdrawal thresholds (MWT) for Arch-K14 Cre+ and Arch-K14 Cre-mice, pre- and post-paclitaxel or vehicle initiation; sample size: n = 7-10. (**B**) Fold change in withdrawal thresholds at each experimental day for vehicle treated Arch-K14 Cre+ and paclitaxel treated Arch-K14 Cre+ animals. Attending responses to dynamic brush (**C**) and noxious needle (**D**) hind paw stimuli, 10 days post-paclitaxel initiation, for both Arch-K14 Cre+ and Arch-K14 Cre-groups; n = 4-5. (**E**) Mechanical withdrawal thresholds (MWT) of both hM4Di-K14 Cre+ and hM4Di-K14 Cre-mice, assessed before (baseline) and following paclitaxel or vehicle administration; n = 9-12. (**F**) Fold change in withdrawal thresholds at each experimental day for vehicle treated hM4Di-K14 Cre+ and paclitaxel treated hM4Di-K14 Cre+ animals. (**G**) Attending responses to dynamic brush (on the left) and noxious needle (on the right) hindpaw stimulation, gauged 10 days post-paclitaxel administration, for hM4Di-K14 Cre+ and hM4Di-K14 Cre-cohorts; n = 9-12. Statistical analysis for (**A, C, D, E, G**, and **H**) was performed using a 3 way ANOVA with Tukey’s multiple comparisons tests; for (**B** and **F**) using a 2 way ANOVA with Tukey’s multiple comparisons tests *P < 0.05, **P < 0.01, ***P < 0.001, and ****P <0.0001. Error bars represent SE. For (**A, C, D, E, G**, and **H**) Black asterisks represent comparisons between vehicle treated animals while red or blue asterisks represent comparisons between paclitaxel treated animals. Schematic created with BioRender.com.

To confirm that keratinocyte inhibition decreases paclitaxel-induced hypersensitivity, we used a complimentary chemogenetic approach. We generated a mouse line that expresses citrine-tagged human muscarinic M4 Designer Receptor Exclusively Activated by Designer Drugs (hM4Di-DREADD) specifically in K14-expressing epidermal cells (hM4Di-K14Cre+ mice) and activated hM4Di using deschloroclozapine (DCZ), a highly selective agonist for muscarinic-based DREADDs (**Fig. S1A**)^13^. Bath application of 0.5 µM of DCZ resulted in membrane potential hyperpolarization of keratinocytes isolated from hM4Di-K14Cre+ mice but not from hM4Di-K14Cre-controls (**Fig. S1D**). Consistent with our optogenetic data, we found that chemogenetic inhibition of keratinocytes increased the von Frey mechanical withdrawal thresholds of paclitaxel-treated hM4Di-K14 Cre+ mice but had no effect on hM4Di-K14 Cre-control mice (**Fig. 1E**). The overall effect of keratinocyte chemogenetic inhibition on mechanical withdrawal thresholds was greater in paclitaxel-treated hM4Di-K14Cre+ mice than in vehicle-treated hM4Di-K14Cre+ mice (**Fig. 1F**). Moreover, chemogenetic inhibition of keratinocytes decreased the paclitaxel-induced mechanical hypersensitivity to both dynamic paintbrush and noxious pinprick stimuli (**Fig. 1G, H**). Our results confirm via two methods that hyperpolarization of keratinocytes decreases paclitaxel-induced mechanical hypersensitivity^8,9^. In summary, these results indicate that keratinocyte activity is required to maintain the persistent paclitaxel-induced mechanical hypersensitivity.

### Keratinocytes in paclitaxel-treated mice are sensitized to mechanical stimulation

Since keratinocyte inhibition decreased paclitaxel induced behavioral mechanical hypersensitivity, we next asked whether paclitaxel treatment sensitizes keratinocytes to mechanical stimulation. To test this, we established a mouse line expressing the genetically encoded calcium indicator GCaMP6 in K14-expressing epidermal cells (GCaMP6-K14Cre+), then used *in vivo* confocal imaging to measure mechanically induced calcium flux in earlobe keratinocytes^14^ (**Fig. 2A**). Keratinocytes in paclitaxel-treated GCaMP6-K14Cre+ mice displayed increased calcium responses upon mechanical stimulation compared to keratinocytes in vehicle-treated mice (**Fig. 2B, C).** Since paclitaxel related pain can especially affect the hands and feet, we tested whether paclitaxel affects the mechanical sensitivity of glabrous hind paw skin keratinocytes. The dense stratum corneum in the mouse paw skin made visualizing keratinocytes in GCaMP6-K14+ mice challenging in *in vivo* settings. Therefore, we conducted experiments on whole epidermis isolated from the glabrous hind paw skin of GCaMP6-K14+ mice treated *in vivo* with paclitaxel (**Fig. 2D**). Consistent with our findings in ear skin, glabrous hindpaw keratinocytes from paclitaxel-treated mice exhibited increased calcium responses to mechanical stimulation (**Fig 2E, F**). Together these results demonstrate that paclitaxel treatment sensitizes keratinocytes to mechanical stimulation *in vivo* and *in vitro* across several body regions. Thus, paclitaxel treatment *in vivo* may affect epidermal keratinocyte function throughout the organism.

**Fig. 2.**
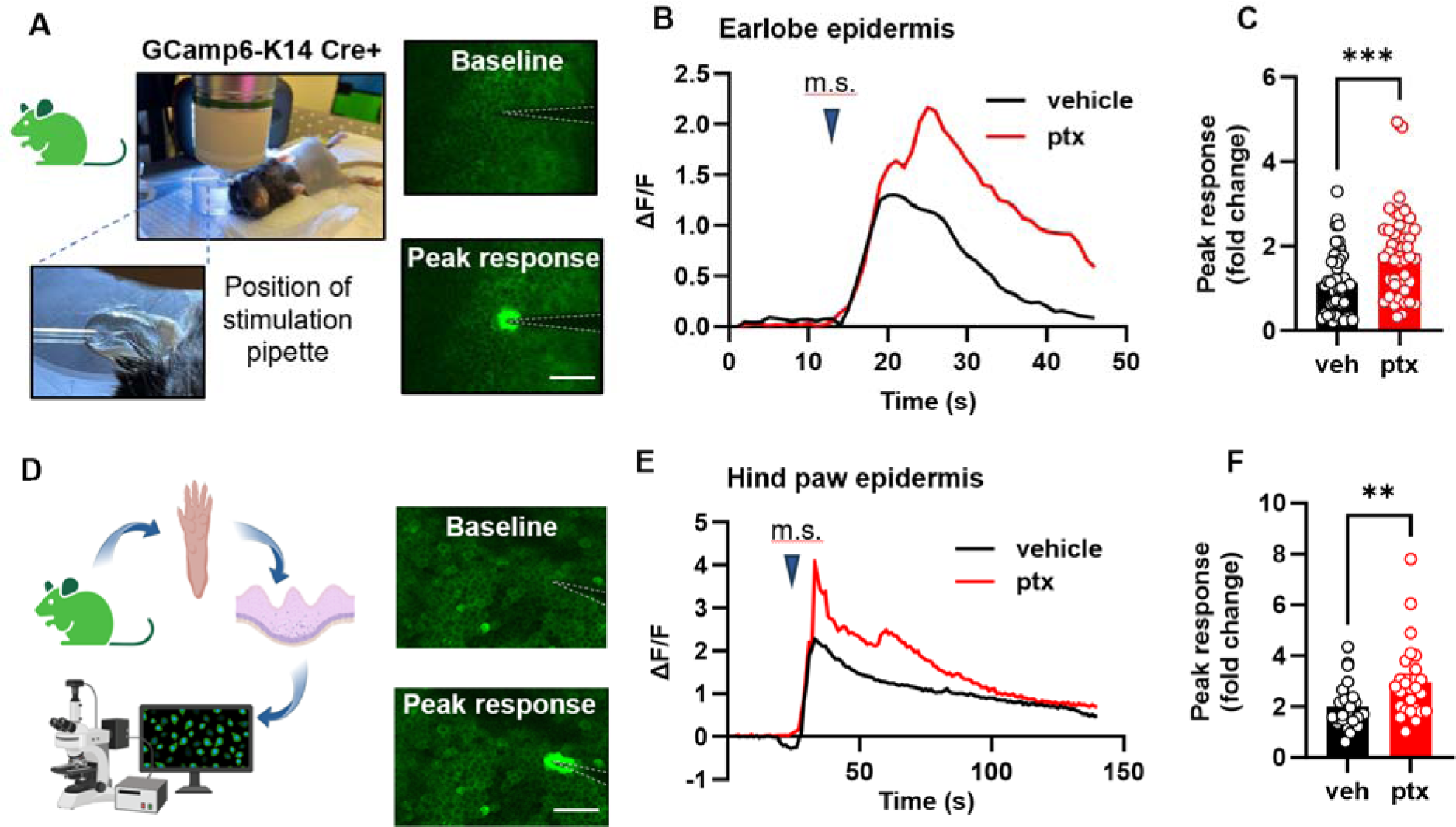
Paclitaxel treatment sensitizes the mouse epidermis to mechanical stimulation. (**A**) Visual representation and example fluorescent images of the in *vivo* mechanical calcium imaging setup used on the earlobe skin of GCamp6-K14 Cre+ mice. (**B**) Example traces illustrating calcium responses induced by mechanical stimulation (m.s.) in earlobe skin. (**C**) Comparative analysis of peak calcium responses to mechanical stimuli of earlobe skin in paclitaxel (ptx) and vehicle-treated mice. Data was sourced from 4 mice per group, with 5-15 recordings per mouse (totaling 45-46 recordings per treatment group). (**D**) Diagram showcasing the experimental approach for *in situ* calcium imaging on paw keratinocytes. Hindpaw skin from GCamp6-K14 Cre+ mice underwent dissection, with the epidermis subsequently separated from the dermis for imaging purposes (left). Fluorescence images of the paw epidermis both at baseline and immediately following mechanical stimulation (right). (**E**) Representative traces displaying calcium responses induced by mechanical stimulation (m.s.) in the hind paw epidermis. (**F**) Comparative analysis of peak calcium responses to mechanical stimuli of hind paw epidermis of paclitaxel (ptx) and vehicle-treated mice (right). Data comprises recordings from 5 mice per group, with 5 recordings taken per mouse (yielding 25 recordings for each treatment group). Scales are 50 µm. Statistical analysis was performed using Student’s T test; **P < 0.01, and ***P < 0.001. Error bars represent SE. Schematic created with BioRender.com.

Next, we asked whether paclitaxel sensitization of keratinocytes persists when the keratinocytes are removed from their native tissue. We isolated keratinocytes from the hind paws of paclitaxel and vehicle treated C57BL/6J mice and measured their mechanical sensitivity using calcium imaging and patch-clamp electrophysiology (**Fig. 3A**). Keratinocytes were mechanically stimulated via a blunted glass probe controlled by a Piezo1 actuator. Stimulation was applied in stepwise increments of 1 μm until the first calcium response was triggered (**Fig. 3B**). Paclitaxel did not affect baseline calcium levels (**Fig. 3C**). Keratinocytes from paclitaxel treated animals displayed decreased mechanical thresholds and increased calcium response magnitudes (**Fig. 3D, E**). In whole-cell patch clamp studies, the mechanically activated current was evoked from individual keratinocytes by a stepwise increase in membrane indentation using a blunt glass probe (**Fig. 3F**). Paclitaxel treatment did not alter the proportions of keratinocytes that responded to mechanical probing with rapidly adapting (RA), intermediately adapting (IA), or slowly adapting (SA) currents (**Fig. 3G**). Keratinocytes from paclitaxel-treated animals did however exhibit reduced mechanical thresholds compared to cells from vehicle-treated controls (**Fig. 3H**); mechanical current density was not affected by paclitaxel (**Fig. 3I**). Together, these findings show that paclitaxel treatment *in vivo* results in mechanical sensitization of isolated keratinocytes in the form of decreased mechanical thresholds and increased calcium response magnitudes.

**Fig. 3.**
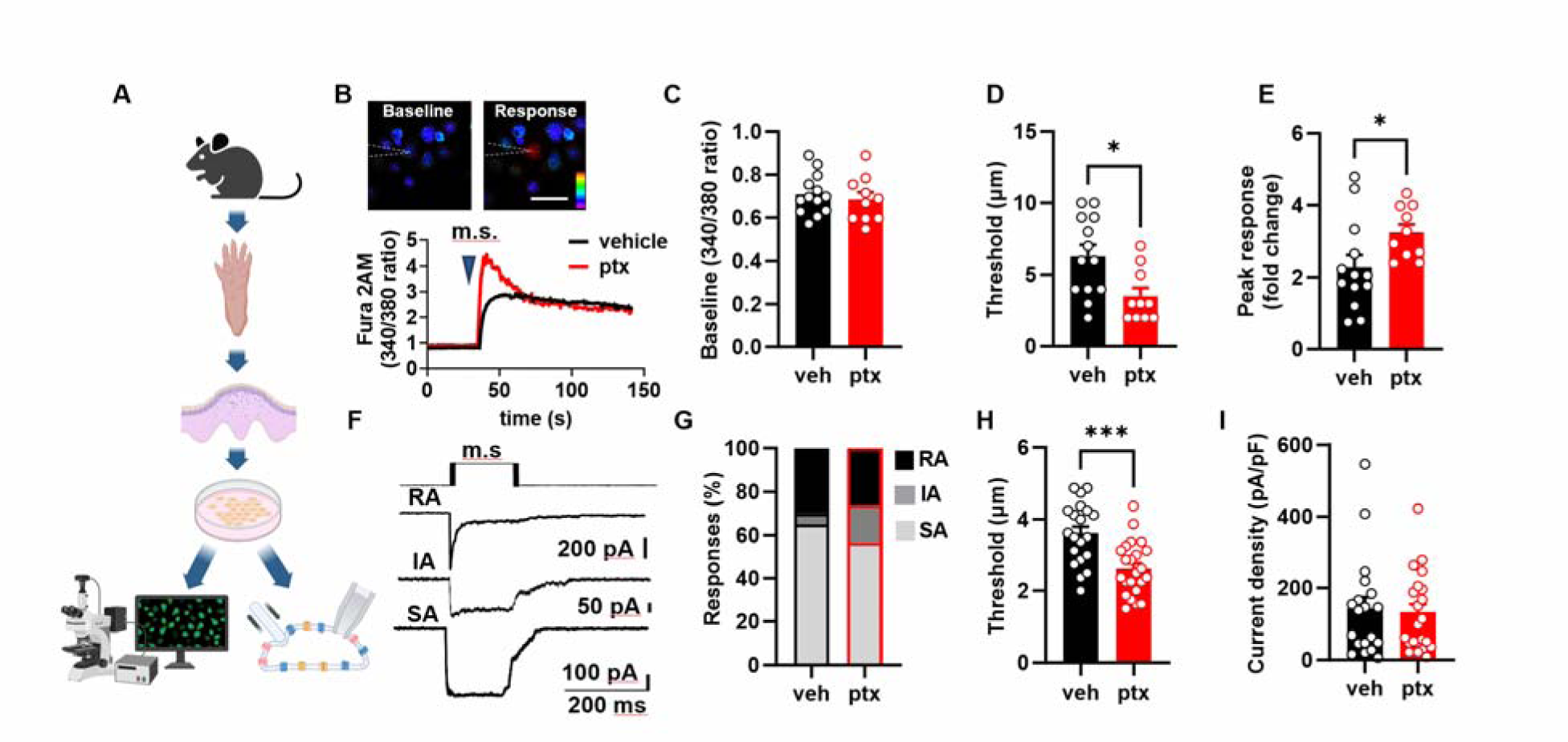
Paclitaxel treatment lowers the threshold of keratinocyte activation by mechanical stimulation: (**A**) Diagram illustrating the procedure to isolate keratinocytes from the mouse hindpaw epidermis for subsequent calcium imaging and patch clamp electrophysiology studies. (**B**) Representative traces accompanied by images illustrating mechanically activated calcium responses. The position of the stimulation electrode is depicted with dashed lines. The time of mechanical stimulation (m.s) is shown as a triangle. The scale is 20 µm. (**C**) Baseline 340/380 fluorescence ratio of keratinocytes derived from mice treated with either paclitaxel (ptx) or vehicle. (**D**) Activation threshold for the mechanically induced calcium responses. (**E**) Amplitude of mechanically activated calcium responses, normalized to baseline 340/380 ratio. (**F**) Example traces showcasing the variations in adaptation rates (rapidly adapting (RA), intermediately adapting (IA), and slowly adapting (SA)) of whole-cell currents induced by membrane indentation (schematic protocol of mechanical stimulation (m.s) is presented on **A**). Comparative analysis of the distribution of different types (**G**), activation threshold (**H**), and density (**I**) of mechanically-activated currents recorded from keratinocytes isolated from vehicle and paclitaxel-treated mice. Statistical analysis was performed using Student’s *t*-test; For (**G**) a chi-square test with Fischer’s Exact post-hoc test was performed *P < 0.05, and ***P < 0.001. Error bars represent SE. Schematic created with BioRender.com.

### Paclitaxel sensitizes Piezo1 in mouse keratinocytes

We recently showed that the mechanically sensitive channel Piezo1 sets the activation threshold of mechanically activated currents in keratinocytes^11^. To determine whether paclitaxel-induced mechanical sensitization of keratinocytes is mediated through Piezo1, we isolated hind paw keratinocytes from paclitaxel or vehicle treated mice and performed calcium imaging with the Piezo1 agonist Yoda1^15^ (**Fig. 4A**). We found that paclitaxel treatment did not increase the number of Yoda1-responsive keratinocytes (**Fig. 4B**). However, the magnitude of Yoda1 induced calcium response was larger in the paclitaxel-treated animals, suggesting that *in vivo* paclitaxel treatment sensitizes keratinocyte Piezo1 current properties (**Fig. 4C**). To determine if keratinocyte Piezo1 sensitization mediates paclitaxel-induced behavioral mechanical hypersensitivity, we generated an epidermal-specific Piezo1 knockout mice (Piezo1-K14Cre+)^11^. Paclitaxel treatment decreased withdrawal thresholds in both Piezo1-K14Cre+ and Piezo1-K14Cre-mice, however, Piezo1-K14Cre+ mice exhibited significantly less mechanical hypersensitivity, which persisted for at least 21 days after the start of paclitaxel treatment (**Fig. 4D**). Moreover, paclitaxel-treated Piezo1-K14Cre+ mice exhibited fewer attending responses in dynamic brush and noxious pinprick assays compared to paclitaxel-treated Piezo1-K14Cre-mice (**Fig. 4E, F**). These results indicate that paclitaxel-induced mechanical hypersensitivity is mediated in part by keratinocyte Piezo1.

**Fig. 4.**
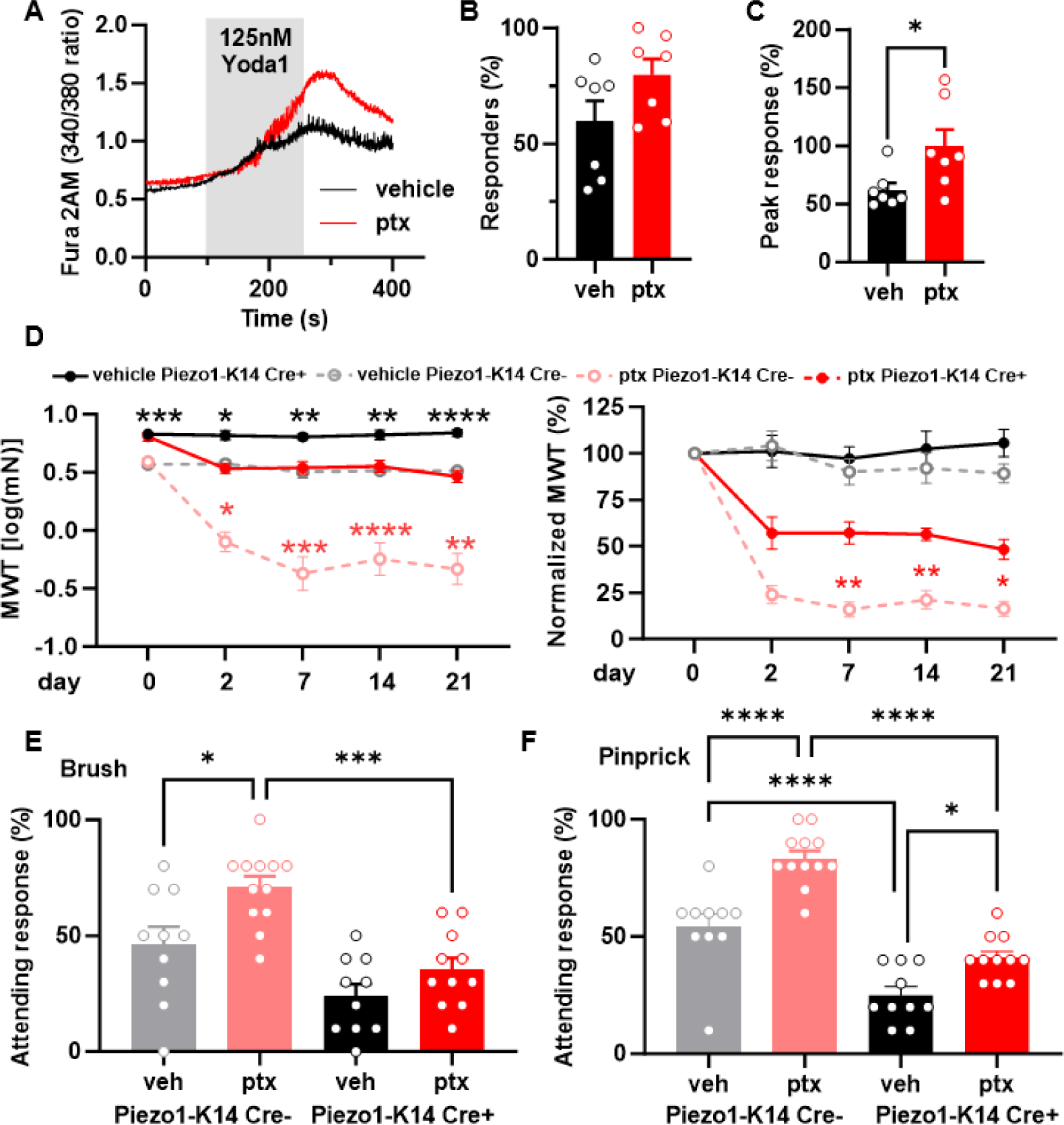
Epidermal Piezo1 mediates paclitaxel-induced mechanical hypersensitivity. (**A**) Representative traces showcasing calcium responses evoked by Yoda1 (125 nM) in keratinocytes derived from either vehicle or paclitaxel-treated mice. (**B**) The proportion of keratinocytes exhibiting a response to Yoda1. Dots depict the average count of responsive keratinocytes per mouse. (**C**) Magnitude of the calcium response to Yoda1. Dots depict the average peak calcium response observed in keratinocytes for each animal (n = 7, 60-100 cells per animal). (**D**) Comparison of mechanical withdrawal thresholds between Piezo1-K14 Cre+ and Piezo1-K14 Cre-mice, both before and after paclitaxel or vehicle treatment (left). Mechanical withdrawal threshold data normalized to baseline values (right); data sourced from 9-12 mice. Evaluation of responses to dynamic brush (**E**) and to the noxious needle (pinprick, **F**) hindpaw stimuli, at day 10 post-paclitaxel injection, in both Piezo1-K14 Cre+ and Piezo1-K14 Cre-groups; sample size: n = 9-12. Statistical analysis for (**B and C**) was performed using a Student’s T test; for (**D**) using a 3 way ANOVA with Tukey’s multiple comparisons test; for (**E and F**) using a 2 way Anova with Tukey’s multiple comparisons test. *P < 0.05, **P < 0.01, ***P < 0.001, and ****P< 0.0001. Error bars represent SE. For figure (**D**), black asterisks represent comparisons between vehicle treated animals while red asterisks represent comparisons between paclitaxel treated animals. Schematic created with BioRender.com.

### Paclitaxel directly sensitizes Piezo1 to mechanical activation

To determine the mechanism through which paclitaxel treatment leads to enhanced keratinocyte Piezo1 responses, we examined if paclitaxel treatment alters Piezo1 expression in keratinocytes. Piezo1 expression was not different between paclitaxel and vehicle treated mouse epidermis, suggesting that the enhanced keratinocyte Piezo1 responses were due to sensitization of Piezo1 rather than increased expression (**Fig. 5A**). To determine whether paclitaxel can directly sensitize Piezo1 channels, we used HEK-P1KO cells (cell line lacking endogenous Piezo1^16^) transfected with eGFP-tagged mouse Piezo1, incubated these cells for 30 minutes with 1 μM paclitaxel or vehicle, and recorded stretch-induced and spontaneous single channel Piezo1 currents ^17,18^. Paclitaxel-treated cells exhibited increased stretch-activated Piezo1 currents compared to vehicle controls (**Fig. 5B-D**). Analysis of spontaneous single-channel Piezo1 demonstrated that paclitaxel treatment significantly increases the unitary amplitude of the Piezo1 channels but does not affect the number of open channels or their open probability (**Fig. 5E-H**). Furthermore, paclitaxel exposure resulted in a significant increase in the single channel conductance of Piezo1 compared to vehicle controls (**Fig. 5I**).

**Fig. 5.**
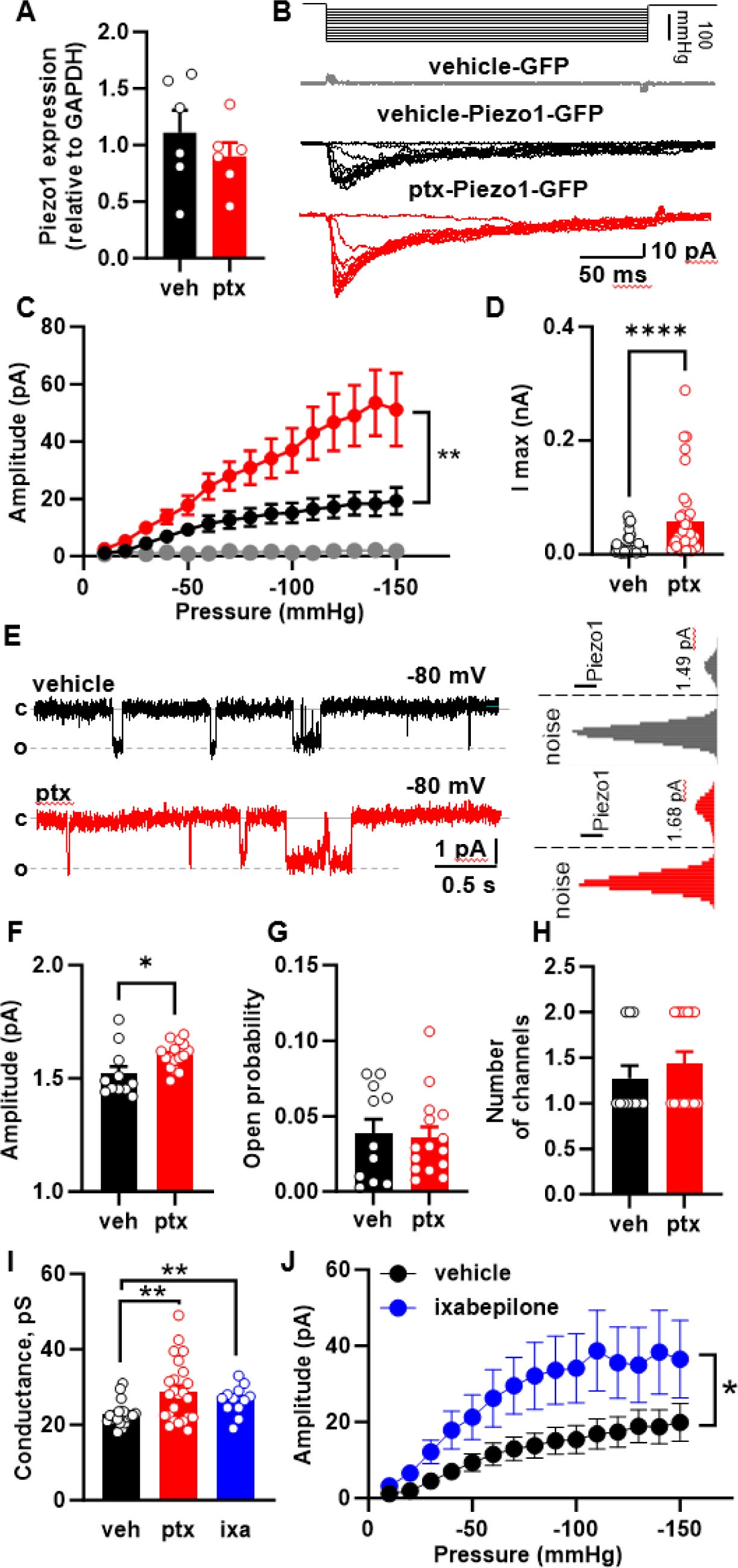
Effect of microtubule-stabilizing drugs on Piezo1 expression and activity. (A) Piezo1 expression in epidermis of mice treated with paclitaxel or vehicle. **(B)** Representative cell-attached recordings of stretch-activated currents induced by a stepwise increase of negative pressure in the patch pipette at V_hold_ = -80 mV. Recordings were made from HEK-P1KO cells expressing GFP (grey –vehicle) and Piezo1-GFP in vehicle (black), and paclitaxel (red) pretreated cells. **(C)** Current-pressure relationship of stretch-activated currents recorded from HEK-P1KO cells expressing GFP (grey, n=6) and Piezo1-GFP in vehicle-(black, n=36) and paclitaxel treated (red, n=39) cells at a V_hold_ = -80 mV. (**D**) The maximal amplitude of stretch-activated Piezo1 current recorded in the vehicle (black, n = 36), and paclitaxel (red, n=39) pretreated HEK-P1KO cells. (**E**) Representative cell-attached recordings of spontaneous single-channel Piezo1 activity in the vehicle- and paclitaxel-treated HEK-P1KO cells with corresponding all-point amplitude histograms. (**F-H**) Summary graphs of single-channel Piezo1 amplitude, open probability, and number of channels recorded from vehicle- and paclitaxel-treated HEK-P1KO cells at V_hold_= -80 mV. (**I**) Effect of paclitaxel and ixabepilone treatment on Piezo1 conductance. (**J**) Current-pressure relationship of stretch-activated currents recorded from Piezo1 expressing HEK-P1KO cells treated with vehicle- (black, n=36) or ixabepilone (1 µM, blue=20). Recordings were made at a V_hold_ = - 80 mV. Statistical analysis for (**C** and **J**) was performed using 2way ANOVA with Tukey’s multiple comparisons tests; for (**D** and **H**) using the Mann-Whitney test, and for (**A, F, G** and **I**) using Student’s *t* test. Error bars, ± SEM. *P < 0.05, **P < 0.01, and ****P < 0.0001.

Paclitaxel is a chemotherapeutic due to its ability to stop tumor proliferation, a task it completes by stabilizing microtubules and reducing cytoskeletal dynamics^19^. Recent research has highlighted the role of the cytoskeleton in the mechanical sensitivity of both Piezo1 and Piezo2^20^. Therefore, we hypothesized that paclitaxel-induced sensitization of Piezo1 may result from microtubule stabilization. To test this hypothesis, we used two different microtubule destabilizing agents, nocodazole and colchicine ^21,22^ to determine if these compounds could decrease paclitaxel-induced Piezo1 sensitization. Neither nocodazole nor colchicine reversed the paclitaxel-induced sensitization of Piezo1 currents (**Fig. S2A-C**). Next, we asked whether a non-taxane microtubule stabilizing agent (ixabepilone), could enhance Piezo1 currents^23^. HEK-P1KO cells transfected with Piezo1 were incubated with ixabepilone for 30 minutes. Similar to the effect of paclitaxel treatment, ixabepilone increased Piezo1 spontaneous single channel conductance and sensitized Piezo1 expressing cells to mechanical stimulation (**Fig 5I, J**). Together these findings suggest that microtubule stabilizers directly sensitize Piezo1 channels. However, microtubule destabilizers were not sufficient to reverse this sensitization once established.

### Paclitaxel sensitizes human keratinocytes to mechanical stimulation and Yoda1 activation

To determine whether our findings in mouse translate to human, we isolated keratinocytes from breast skin of healthy donors. Human keratinocytes were exposed to paclitaxel (1 μM) or vehicle for 30 minutes (**Fig. 6A**). Human keratinocytes exposed to paclitaxel exhibited increased mechanically induced calcium responses compared to vehicle treated cells (**Fig.6B, C**). The mechanical threshold that initiated this response and the baseline calcium levels were unaffected by paclitaxel exposure (**Fig. 6D, E**). We hypothesized that the paclitaxel-induced mechanical sensitization was mediated by Piezo1^11,24^. Supporting this, paclitaxel pretreatment sensitized human keratinocytes to Yoda1 activation, increasing both the number of keratinocytes responding and the magnitude of calcium influx elicited by Yoda1 (**Fig. 6F-H**). Similarly, ixabepilone, a non-taxane microtubule stabilizer, sensitized human keratinocytes to Yoda1 application (**Fig. 6I, J**). Together, these data indicate that microtubule stabilizers directly sensitize human keratinocytes to mechanical stimulation and Piezo1 activation.

**Fig. 6.**
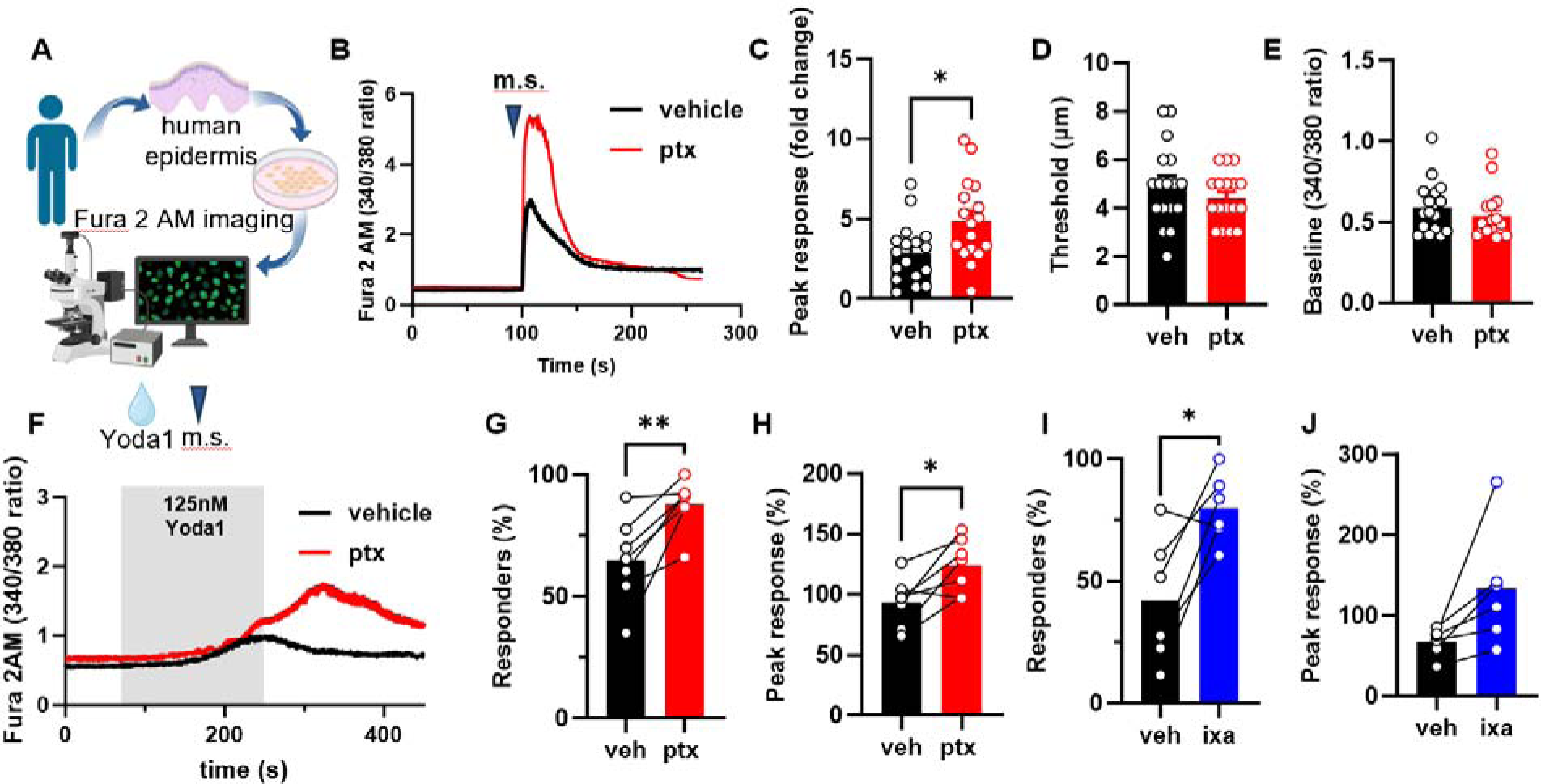
Human keratinocytes are sensitized to mechanical stimulation and Yoda1 activation following direct paclitaxel and ixabepilone exposure. (**A**) Schematic of human keratinocyte isolation from tissue donor samples and subsequent Fura 2 calcium imaging. (**B**) Representative traces of calcium responses evoked by mechanical stimulation (m.s.) in paclitaxel or vehicle-treated human keratinocytes (keratinocytes were treated with 1 μM paclitaxel or vehicle for 30 minutes before imaging). (**C**) Amplitude of mechanically activated calcium responses. (**D**) Activation threshold for the mechanically induced calcium responses. (**E**) Baseline 340/380 fluorescence ratio recorded from human keratinocytes incubated with paclitaxel or vehicle. (**F**) Example traces of Yoda1 (125 nM) evoked calcium responses in human paclitaxel or vehicle-treated keratinocytes. The time of Yoda1 application is depicted with a grey box. (**G**) Proportion of human keratinocytes treated with paclitaxel or vehicle exhibiting a response to Yoda1. (**H**) Magnitude of the calcium response to Yoda1 in human keratinocytes treated with paclitaxel or vehicle (right). (**I**) Proportion of human keratinocytes treated with ixabepilone or vehicle exhibiting a response to Yoda1. (**J**) Magnitude of the calcium response to Yoda1 in human keratinocytes treated with ixabepilone or vehicle. Dots depict the average peak calcium response observed in keratinocytes for each human sample (n = 6-7, 25-120 cells per n). Connecting lines depict cells from the same human donor in two different treatment conditions. Statistical analysis for (**C-E**) was performed using a Student’s t test, and for (**G-J**) using a paired Student’s t test. Error bars, ± SEM. *P < 0.05, **P < 0.01, and ****P < 0.0001.

## DISCUSSION

Over the last decade, it has become clear that non-neuronal cutaneous cells play an essential role in sensing exogenous innocuous and noxious mechanical and thermal stimuli applied to healthy skin^8,9,11,25–28^. This discovery has led to a shift in our understanding of somatosensation because the peripheral terminals of sensory neurons were long considered the primary transducers of sensory stimuli in the skin. However, whether non-neuronal skin cells contribute to the pain and hypersensitivity that develops after skin injury or disease of any type is unknown. Here, we investigated the contribution of epidermal keratinocytes to the mechanical hypersensitivity induced by paclitaxel. Our findings reveal for the first time that keratinocytes are key drivers of mechanical pain following chemotherapy treatment.

We found that either optogenetic or chemogenetic inhibition of keratinocytes markedly attenuated both innocuous and noxious touch hypersensitivity in paclitaxel treated mice, from as early as 2-days to at 3-weeks post paclitaxel treatment. These results suggest keratinocyte signaling is required to maintain paclitaxel-induced mechanical hypersensitivity. Notably, keratinocyte signaling appears to be fast; the withdrawal thresholds of archaerhodopsin expressing mice were increased within 1-minute of laser stimulation and returned to pre-laser withdrawal thresholds within 1-minute post-laser stimulation. This keratinocyte signaling may be mediated through mechanisms such as gap junctions or synaptic signaling with sensory nerve fibers^12^. Sensory nerve terminals have been shown to be enwrapped within keratinocyte cytoplasmic gutters as they innervate the skin. Indeed, skin biopsies from patients with small-fiber neuropathy show increased numbers of synaptic vesicles present in keratinocytes adjacent to sensory nerve endings, suggesting the cellular machinery is/morphological structures are present for amplified keratinocyte signaling following injury^12^. It is important to note that neither keratinocyte optogenetic nor chemogenetic inhibition completely reversed the mechanical hypersensitivity induced by paclitaxel. This residual hypersensitivity is likely mediated by the actions of paclitaxel on dorsal root ganglion and spinal cord nociceptors^29^. An important question remains regarding the identity of the signaling mediators and mechanisms that underly keratinocyte to sensory nerve communication. Prior studies established that mechanical stimulation induces the release of ATP from keratinocytes^9,30–32^, a mechanism that is essential for normal behavioral responses to touch stimuli in mice^9,28^. This release of ATP from keratinocytes can be reduced by membrane hyperpolarization, which both Archaerhodopsin and hM4Di activation achieve^8,9,33,34^. Interestingly, antagonists of the ATP receptors P2×7 and P2×4 reduce paclitaxel-induced mechanical hypersensitivity^35^. Thus, increased keratinocyte to sensory neuron purinergic signaling may mediate all or part of the paclitaxel-induced mechanical hypersensitivity. Beyond ATP, keratinocytes release a variety of neuroactive factors, including glutamate, CGRPβ, β-endorphins, and endothelin-1, which can act directly on sensory fibers^36–40^. Future studies should explore the specific molecules and mechanisms underlying keratinocyte to sensory nerve signaling and how these may change following paclitaxel hypersensitivity and other types of tissue injury.

We found that keratinocytes, from mice treated with paclitaxel and human keratinocytes directly exposed to paclitaxel, exhibit increased sensitivity to mechanical stimulation. This heightened sensitivity involves the mechanotransducer Piezo1, as paclitaxel treated mouse and human keratinocytes were sensitized to Yoda1 activation. Notably, deleting Piezo1 from keratinocytes partially protected against paclitaxel-induced behavioral mechanical hypersensitivity. Together, these findings suggest that amplified mechanical responsiveness of keratinocytes and sensitization of Piezo1 plays a significant role in paclitaxel-induced mechanical hypersensitivity. Furthermore, the sensitization of human keratinocytes by paclitaxel exposure suggests that our findings in mice may be translatable to human cases of paclitaxel-induced mechanical hypersensitivity.

Our results indicate that paclitaxel exposure is sufficient to sensitize Piezo1. This effect appears to stem from heightened Piezo1 channel sensitivity rather than an increase in Piezo1 expression. Indeed, paclitaxel has previously been shown to increase the conductance of TRPA1^41^. Similarly, our findings suggest that brief paclitaxel exposure (30 minutes) can modulate Piezo1 single channel conductance and mechanical sensitivity. Given paclitaxel’s role as a microtubule stabilizer, we hypothesized that this stabilization could be driving Piezo1 sensitization, considering the intimate connection between the cytoskeleton and mechanically activated channels^20^. Testing with ixabepilone, a non-taxane microtubule stabilizer, similarly enhanced the sensitivity of Piezo1-expressing HEK cells to mechanical stimulation and human keratinocytes to Yoda1 activation. This suggests microtubule stabilization’s involvement in paclitaxel-induced Piezo1 sensitization. However, attempts to reverse the effects of paclitaxel with microtubule destabilizers, nocodazole and colchicine, proved unsuccessful. This may be due to these microtubule destabilizers’ inability to sufficiently block or reverse paclitaxel’s effects when applied together. However, alternate paclitaxel-induced Piezo1 sensitization mechanisms – such as channel post-translational modifications or membrane lipid composition changes – cannot be dismissed and should be explored in future studies^42,43^. Paclitaxel’s ability to sensitize Piezo1 could have broader repercussions beyond pain. Given the role of keratinocyte Piezo1 in epidermal function, enhanced Piezo1 activity might contribute to the adverse skin reactions and poor wound healing associated with paclitaxel treatment^44–46^. Moreover, its expression and sensitization across different tissues could be linked to other side effects of body-wide paclitaxel treatment, including cardiac issues, respiratory complications, urinary problems, and anemia^2,47–50^. Future research should address the potential role of paclitaxel-induced Piezo1 sensitization in these contexts.

While keratinocytes are the predominant K14 expressing cells in the epidermis, Merkel cells, also K14 expressing, are important for light touch detection^25,51,52^. These cells detect fine mechanical forces via connections with Aβ slowly adapting Type I low threshold mechanoreceptors ^53–55^. Aβ mechanoreceptor activity is required for paclitaxel-induced mechanical allodynia and previous research indicates that paclitaxel exposure heightens Merkel cell responses to mechanical stimuli^56,57^. Thus, in our optogenetic/chemogenetic behavioral experiments, we cannot rule out the potential contribution of Merkel cells to paclitaxel-induced mechanical hypersensitivity. Although Merkel cells form synapses with these mechanoreceptors, keratinocytes are also close to these fibers in the basal epidermis^51,52,58^. While direct interaction of keratinocytes with Aβ mechanoreceptors remains uncertain, genetic ablation of Merkel cells showed minimal impact on mechanical allodynia in a spared nerve injury model of neuropathic pain^59^. Furthermore, diminished responses to mechanical stimulation following Piezo1 deletion in the epidermis likely reflect reduced keratinocyte mechanical signaling, as Merkel cell sensitivity primarily depends on Piezo2^11,51^. Importantly, Merkel cells are unlikely to contribute to noxious needle stimulation responses, which are mediated by A-fiber nociceptors^60,61^. Overall, our findings demonstrate a significant role of epidermal cells in paclitaxel-induced mechanical hypersensitivity. Paclitaxel-induced hypersensitivity to light touch may be driven by signaling from keratinocytes and/or Merkel cells to low threshold sensory afferents, while hypersensitivity to noxious touch stimuli is mediated by increased keratinocyte signaling to high threshold sensory afferents. Further research is needed to clearly define the roles of keratinocytes and Merkel cells in mechanical hypersensitivity.

This is the first study to demonstrate the role for keratinocyte mechanotransduction and Piezo1 in the maintenance of injury-induced mechanical hypersensitivity. Our findings provide rational for the development of therapeutics targeted at keratinocytes and epidermal Piezo1 for alleviating paclitaxel-induced pain. Furthermore, alterations in keratinocyte function and signaling have been described in a variety of painful inflammatory and neuropathic conditions that affect the skin; these include the devastating painful conditions complex regional pain syndrome, post-herpetic neuralgia, and fibromyalgia^62,63^. The potential contribution of keratinocyte mechanotransduction to mechanical pain in these conditions should be explored.

## MATERIALS AND METHODS

### Study Design

This study aimed to assess the extent to which keratinocytes and epidermal Piezo1 contribute to persistent mechanical pain in chemotherapy-induced peripheral neuropathy. To achieve this, we performed reflexive behavioral tests of mechanical sensitivity in a rodent model of paclitaxel-induced neuropathic pain using mouse lines that express inhibitory optogenetic (Archaerhodopsin) and chemogenetic proteins (hM4Di) specifically in the epidermis. We employed confocal calcium imaging microscopy and transgenic mice which selectively express calcium indicator GCamp6 in keratinocytes to evaluate the effect of paclitaxel treatment on the mechanical sensitivity of mouse earlobe and glabrous hind paw epidermis *in vivo* and *in situ*. Patch clamp electrophysiology was used to evaluate the mechanosensitivity of mouse keratinocytes *in vitro*. To model the effects of paclitaxel treatment in the human epidermis, we isolated keratinocytes from the skin of human tissue donors and pretreated them with paclitaxel (or ixabepilone). Fluorescence calcium imaging microscopy was employed to evaluate a single keratinocyte’s mechanical sensitivity and alteration of Piezo1 signaling in mouse and human keratinocytes following paclitaxel treatment. To determine the extent to which epidermal Piezo1 contributes to paclitaxel-induced mechanical hypersensitivity we performed reflexive behavior assays in transgenic mice with a selective deletion of PIEZO1 in keratinocytes. To determine the mechanisms of Piezo1 sensitization by paclitaxel, we pretreated Piezo1 expressing HEK293 cells directly with paclitaxel or vehicle and performed electrophysiological recordings measuring spontaneous and stretch activated Piezo1 activity. Sample sizes were determined based on previous behavioral and electrophysiological data published by our laboratory. In all experiments, animals and cells were randomly assigned to treatment groups. Experimenters were blinded to treatment and genotype during data acquisition and analysis. The number of animals or cells used is noted in all figure legends.

### Animals

For all experiments, male and female mice of at least 8 weeks of age were used. Animals had ad libitum access to food and water and were housed in a climate-controlled room with a 12:12 light:dark cycle, on Sani-Chips aspen wood chip bedding (P.J. Murphy Forest and Products, New Jersey) with a single pack of ENVIROPAK nesting material (W.F Fisher & Son, Inc, New Jersey). All animals were group housed with a minimum of 3 mice per cage. All protocols were in accordance with the National Institute for Health guidelines and were approved by the institutional Animal Care and Use Committee at the Medical College of Wisconsin (Milwaukee, WI; protocol no. 383).

Adult male and female C57BL/6J mice from Jackson Laboratories (Jackson stock number 000664; Bar Harbor, Maine) were used for Yoda1 calcium imaging and patch-clamp recording studies. A Keratin14(Krt14)Cre driver was used to target epidermal keratinocytes^64–66^. To create mice that express GFP-tagged Archaerhodopsin-3 in K14-positive cells, Ai35D (B6;129S-Gt(ROSA)26Sor^tm35.1(CAG-aop3/GFP)Hze^/J) (Jackson stock number 012735) and B6N.Cg-Tg(KRT14-cre)1Amc (Jackson stock number 018964) lines were mated. The offspring were genotyped as either Arch-K14Cre+ or Arch-K14Cre-littermates (controls). To create mice that express the citrine-tagged inhibitory Gi Designer Receptor Exclusively Activated by Designer Drug (DREADD) hM4Di in K14-positive cells, R26-hM4Di/mCitrine (B6.129-*Gt(ROSA)26Sor^tm1(CAG-CHRM4*,-mCitrine)Ute^*/J) (Jackson stock number 028866) and KRT14-cre lines were mated. Offspring were genotyped as either hM4Di-K14Cre+ or hM4Di-K14 Cre-littermates (controls). To create mice that express the GCaMP6 calcium indicator in K14-positive cells, Ai96 (B6J.Cg-*Gt(ROSA)26Sor^tm96(CAG-GCaMP6s)Hze^*/MwarJ) (Jackson stock number 024106) and KRT14-cre lines were mated. Offspring genotyped as GCaMP6-K14Cre+ were used experimentally. To create mice that lacked Piezo1 in K14-expressing cells, P1^f^ (B6.Cg-*Piezo1^tm2.1Apat^*/J) (Jackson stock number 029213) and KRT14-cre lines were mated. Offspring were genotyped as either Piezo1-K14Cre+ or Piezo1-Krt14Cre^-^ littermates.

### Paclitaxel treatment

Paclitaxel (Taxol, European Pharmacopoeia, Sigma-Aldrich) was dissolved in vehicle (16% ethanol, 16% Cremophor, and 68% saline) to achieve a working concentration of 0.8mg/ml. Mice received intraperitoneal injections of paclitaxel (8mg/kg) or vehicle every other day for a total of 4 injections^67^.

For cellular experiments, paclitaxel was dissolved directly in DMSO to make a 2 mM stock solution. This stock solution was then dissolved in the extracellular buffer to make a working solution of 1 μM paclitaxel and applied directly to cultured cells for 30 minutes prior to electrophysiological or calcium imaging recordings^17,18^.

### Behavioral assays

Animals were tested between 8am and 1pm and were allowed a minimum acclimation period of at least 1 hour to the environment and additional 30 minutes to the experimenter before any tests were conducted. Mechanical sensitivity of the glabrous hindpaw skin was gauged using the Up-Down method, employing a sequence of calibrated von Frey filaments ranging from 0.38-37 mN. A paintbrush was utilized to determine sensitivity to dynamic light touch (Brush test), whereas a spinal needle tip was employed to evaluate sensitivity to noxious mechanical stimuli (Pinprick test). In both assessments, the stimulus was administered to the plantar hindpaw a total of 10 times. Any withdrawal responses accompanied by attending behaviors (prolonged paw elevation, flicking, and licking of the paw) were recorded. Baseline withdrawal thresholds were established prior to paclitaxel treatment and subsequently evaluated at intervals of 2, 7, 14, and 21 days post the initial paclitaxel administration. Both Brush and Pinprick tests were performed 10 days after the first paclitaxel injection.

For optogenetic experiments, an amber laser with a wavelength of 590 nm and a power output of 36 mW (ThorLabs) was used to activate Archaerhodopsin expressing cells. Archaerhodopsin is a proton pump that can be activated by 590nm light leading to cell membrane hyperpolarization^8,9^. One minute before conducting behavioral assessments, the fiber optic cable was positioned approximately 10 cm beneath the plantar surface of the mouse’s hind paw. This alignment ensured the laser directly illuminated the central region of the paw. The hind paw remained consistently exposed to the laser’s emission during each mechanical test.

For chemogenetic experiments, deschloroclozapine (DCZ) (Tocris), a potent agonist of muscarinic DREADD receptors, was used to activate hM4Di+ cells. DCZ was dissolved in 0.9% saline and (100 μg/kg) was administered intraperitonially to mice 30 minutes prior to behavioral testing.

### Primary mouse and human keratinocyte cell culture

Mouse glabrous hindpaw skin was dissected and incubated at room temperature in 10 mg/mL dispase (Gibco, ThermoFisher Scientific, Waltham, MA) dissolved in Hank’s Balanced Salt Solution (HBSS) (Gibco) for 45 minutes. Human female breast tissue skin (age range 19-80) was procured from tissue biopsies through the MCW tissue bank. Human skin was incubated overnight at 4 ° C in 10 mg/mL dispase. Following incubation with dispase, the epidermal sheet was separated from the dermis using forceps and incubated at room temperature for 27 min in 50 % EDTA (Sigma-Aldrich) and 0.05% trypsin (Sigma) dissolved in HBSS (Gibco). After incubation with trypsin, heat inactivated fetal bovine serum (Thermo Fisher Scientific) was added and the epidermal sheets were rubbed against the base of a petri dish with a pipette tip to isolate individual keratinocytes. Keratinocytes were plated on laminin coated glass coverslips and allowed to grow for 3 days in EpiLife media (Gibco) with 1% human keratinocyte growth supplement, 0.2% Gibco^TM^ Amphotericin B (250 µg/mL of Amphotericin B and 205 µg/mL sodium deoxycholate, Gibco) and 0.25% penicillin-streptomycin (Gibco). Cultures were kept at 37 °C and 5% CO_2_ conditions. Growth media was exchanged 48 hr after plating.

### RNA isolation and quantitative real-time PCR

The glabrous hindpaw skin of paclitaxel and vehicle treated mice (7 days post first paclitaxel injection) was dissected and the epidermis was separated from the dermis (as described above in *primary keratinocyte culture*). Tissue was homogenized using a Bead Mill (VWR) and lysis buffer containing 1% 2-mercaptoethanol. RNA was isolated using a PureLink RNA Mini Kit (Invitrogen, Life Technologies). Total RNA content was assessed with a Nanodrop Light Spectrophotometer (Thermo Scientific). cDNA was generated using the Superscript VILO cDNA Synthesis Kit (Invitrogen, Life Technologies). The quantitative real-time PCR reaction was performed using a Bio-Rad CFX96 Touch Real-Time PCR Detection System. All samples were run in triplicate and gene expression was normalized to GAPDH. The primer sequences used were as follows: *mPiezo*1-qF: CTTACACGGTTGCTGGTTGG; *mPiezo1*-qR: CACTTGATGAGGGCGGAAT; *mGAPDH*-qF: CCTCGTCCCGTAGACAAAATG; *mGAPDH*-qR: TGAGGTCAATGAAGGGGTCGT.

### Fura2 *in vitro* calcium imaging

Keratinocytes were isolated from the glabrous hind paw skin of paclitaxel and vehicle treated mice 5-8 days after the first injection. For calcium imaging on human keratinocyte, keratinocytes were isolated from human tissue biopsies. Calcium imaging was performed on mouse and human keratinocytes on their third day in culture. Keratinocytes were loaded with 2.5 µg/mL Fura-2AM, a dual-wavelength ratiometric calcium indicator dye, in 2% BSA for 45 min at RT then washed with extracellular buffer (pH 7.4, 320 Osm) containing (in mM) 150 NaCl, 10 HEPES, 8 glucose, 5.6 KCl, 2 CaCl_2_, and 1 MgCl_2_ for 30 min. Keratinocytes were superfused with extracellular buffer, and viewed on a Nikon Eclipse TE200 inverted microscope. Nikon elements (Nikon Instruments, Melville, NY) was used to capture fluorescence images at 340 and 380 nm. For calcium imaging on human keratinocytes, cultured cells were incubated with 1 μM paclitaxel or vehicle 30 minutes prior to imaging. Responsive cells were those that exhibited >30% increase in 340/380 nm ratio from baseline. PIEZO1 agonist, Yoda 1 (125 nM in extracellular buffer) was applied by flow (0.1 ml/s) for 3 min and washed out for 3 min. This concentration of Yoda-1 was chosen based on our previous calcium imaging findings in keratinocytes (Mikesell 2022).

Mechanical stimulation of mouse and human keratinocytes was performed with a blunted glass rod positioned approximately 2 μm from the keratinocyte’s cell membrane and driven by a piezo servo controller (E-625 PZT, Physik Instrumente, Auburn, MA, USA). Keratinocytes were stimulated in a series of mechanical steps in 1 μm increments applied for 150 ms every 10 s. Because of the long decay period of mechanically induced calcium responses, the cell was stimulated until the first mechanically induced calcium response (a response >30% of the baseline 340/380 ratio), after which stimulation was halted and the calcium response recorded.

### *In vivo* and *in situ* mechanical calcium imaging

Adult male and female GCamp6-K14 Cre+ mice were treated with intraperitoneal (i.p.) injections of paclitaxel or vehicle as described above. Intravital imaging of earlobe epidermal cells was performed on day 6-8 after the first injection of paclitaxel/vehicle. Before imaging, mice were anesthetized with i.p. injection of 100 mg/kg ketamine/0.5 mg/kg dexmedetomidine mixture, placed on a heated pad (37°C), and surrounded by warm supportive material to prevent hypothermia and movement while still allowing the mice to breathe easily. The mouse’s outer ear was fixed “ventral portion up” on the plastic platform with double-sided tape. The helix of the ear was cleaned with phosphate-buffered saline (PBS, Gibco), then moisturized with a drop of PBS, and positioned under x20 water immersion objective lens (W Plan-APOCHROMAT 20X, Zeiss, Germany). Mechanical stimuli were manually applied to the surface of the earlobe skin with a blunt glass probe using a water hydraulic micromanipulator (HNW-3; Narishige group, Japan). The glass probe was lowered until brief contact with the earlobe skin was achieved, after which it was retracted to just above the skin. Mechanically evoked calcium signals were recorded from the basal keratinocytes using a spinning disk upright confocal microscope (Intelligent Imaging Innovations, Inc., USA) equipped with a high-speed Orca Flash 4.0 camera (Hamamatsu, Japan). Data was analyzed using ImageJ. At the end of experiments, mice were given i.p. 2.5 mg/kg injections of atipamezole to reverse the effects of dexmedetomidine. For confocal calcium imaging experiments on mouse hind paw epidermis, glabrous skin from hind paw of paclitaxel or vehicle-treated GCamp6-K14Cre+ mice (Day 8 after the first paclitaxel injection) were dissected, and the epidermis was isolated as reported above. For imaging, mouse epidermis was placed “basal layer up” in 60 mm Petri dish filled with ENH and fixed with a slice anchor (SHD-26GH/10, Warner Instruments, Holliston, MA) on the confocal microscope stage. Mechanical stimulation of keratinocytes and calcium imaging from hind paw epidermis was done using the same approach as described for the *in vivo* study above.

### Immunohistochemistry

The glabrous skin of the mouse hindpaw was dissected and cultured as described above. Cultured keratinocytes were then fixed in 4% paraformaldehyde for 20 minutes 2 days after platting. Cells were washed and then blocked for 30 min in SuperBlock™ Blocking Buffer (Thermo) with 0.5% Triton x100, 1.5% NDS, and 0.5% Gelatin. After rinsing, sections were incubated with mouse hosted Cytokeratin 14 (1:500; Abcam ab7800) for 60 min at RT; then incubated for 60 minutes at RT in donkey anti-mouse IgG (H+L) (Alexa Fluor, 647, 2ug/mL) (Invitrogen A-11032). Coverslips were washed, stained with DAPI (Thermo D1306) for 3 min, washed again and then mounted onto coverslips with ProLong™ Gold Antifade Mountant (Thermo P36930). Immunofluorescent images were taken on a Nikon A1 Laser Scanning Confocal Microscope equipped with Nikon NIS-Elements AR v.5.21.03 software. Three animals for each genotype were used.

### Electrophysiology

Keratinocytes were isolated from adult C57BL/6J mice 24 hours after the last injection of paclitaxel or vehicle and cultured for 24-48 hours. Whole-cell recordings were made in voltage-clamp configuration (holding potential -40 mV) with a borosilicate glass pipette (Sutter Instrument Company, Novato, CA) filled with intracellular solution containing (in mM): 135 KCl, 1 MgCl2, 1 EGTA, 0.2 NaGTP, 2.5 ATPNa2, 10 glucose and 10 HEPES (pH 7.25-7.35). Extracellular normal HEPES (ENH) solution contained (in mM): 127 NaCl, 3 KCl, 2.5 CaCl2,1 MgCl2, 10 HEPES, and 10 glucose, pH 7.35-7.40). Mechanical stimulation was elicited using a glass rod positioned in a proximity to the keratinocyte’s membrane and driven by a piezo stack actuator (PA25, PiezoSystem Jena, Jena, Germany) at a speed of 39.17 μm/ms. Keratinocytes were stimulated with a series of mechanical steps in 0.125 μm increments applied for 150 ms every 10 s. Current profiles were classified using the following parameters: rapidly adapting (RA, inactivation time constant (τ) > 10 ms), intermediately adapting (IA, 10 ms < τ < 30 ms), and slowly adapting (SA; τ < 30 ms) current. The effect of the application of Designer Receptors Exclusively Activated by Designer Drug (DREADD) ligand, deschloroclozapine (DCZ) of membrane potential was studied in keratinocytes isolated from six weeks old hM4Di-K14Cre+ or hM4Di-K14Cre-mice and cultured for 24-48 hrs. Current-clamp measurements of membrane potential were performed in tight-seal (seal resistance >2GΩ) cell attach configuration as was previously reported^68,69^. Briefly, keratinocytes were superfused with ENH solution, and membrane potential was recorded across a cell-attached patch at a holding current of zero with patch pipettes filled with ENH solution. Changes in membrane potential values were estimated at 0, 5, 10, and 15 min after bath application of 0.5 µM DCZ. Data was recorded using Axon pCLAMP 11.2 software using Digidata 1550B and Axopatch 200B amplifier and analyzed by Clampfit 11.2 (Molecular Devices LLC, San Jose, CA) and GraphPad Prism 10 software (GraphPad Software, San Diego, CA).

Transient expression of Piezo1 was performed in Piezo1 deficient HEK293T (HEK-P1KO) cell lines to avoid any interference of our recordings with endogenous PIEZO1 channels’ activity (ATCC, CRL-3519 ™)^16^. For expression of PIEZO1 in HEK-P1KO cells, the cells were transiently transfected with green fluorescent protein (eGFP) (0.25 µg of cDNA/ml) and PIEZO1 channel plasmid (Addgene, #80925, 3 µg of cDNA/ml). For control experiments, eGFP was expressed in HEK-P1KO cells without PIEZO1 added. After transfection, cells were cultured on coverslips for 24-48 hours, then treated for 30 min with the vehicle, paclitaxel (1µM), ixabepilone (1µM), or a combination of paclitaxel (1µM) with nocodazole (1 µM) or colchicine (5 µM). The PIEZO1 activity was recorded in a cell-attached mode in a voltage-clamp configuration from GFP-positive cells using patch pipettes filled with a solution of the following composition (in mM): 130 NaCl, 5 KCl, 1CaCl2, 1 MgCl2, 10 TEACl, and 10 HEPES (pH 7.35). To zero membrane potential, we used an extracellular solution of the following composition (in mM): 140 KCl, 1 MgCl2, 10 glucose, and 10 HEPES (pH 7.35)^10^. Data were acquired and subsequently analyzed with an Axopatch 200B amplifier interfaced via a Digidata 1550B to a PC running the pClamp 11.2 suite of software (Molecular Devices, San Jose, CA). The recordings were filtered with an 8-pole, low-pass Bessel filter LPF-8 (Warner Instruments LLC, Hamden, CT) at 0.5 kHz. Stretch-activated currents were elicited by applying brief negative pressure pulses (pulse duration: 250 ms, increments: 10 mmHg, interstimulus interval: 10 s) on membrane patches through the recording electrode using pressure clamp HSPC-1 (ALA Scientific Instruments Inc, Farmingdale, NY). Spontaneous single-channel activity was analyzed over a span of 60–120 s for each experimental condition at a holding potential of -80 mV. Automatically detected event histograms fit with single or multiple Gaussian distributions were used to determine unitary current amplitudes (Clampfit 11.2 software). Channel conductance was evaluated by plotting the amplitudes of the single-channel currents against applied pipette potential, fitting them by linear regression, and estimating slope conductance between -120 and - 40 mV. Channel numbers and open probability (Po) were also assessed using Clampfit 11.2 software. All electrophysiological experiments were performed at room temperature (22-24°C).

### Statistical analysis

Individual data points were presented whenever possible in addition to group means ± SEM. Data was analyzed using GraphPad Prism 10, and results were considered statistically significant when p < 0.05. Specific statistical comparisons used for each data set are indicated in figure legends. Statistical comparisons were performed after consultation with biostatistician Dr. Aniko Szabo. Because of the large number of cells collected per subject in Fura 2 Yoda1 calcium imaging, percent responder data and magnitude of response were analyzed by averaging responses per animal or human donor. For all mouse studies, both male and female animals were used, and data was analyzed independently by sex. No significant differences were noted, so data were reanalyzed with sexes combined and presented together in all graphs. For human samples, only female tissue donors were used.

## ACKNOWLEDGEMENTS

The authors thank the Medical College of Wisconsin Tissue Bank for providing human skin tissues. We also thank the Engineering Core Team at the Medical College of Wisconsin for providing technical support and 3D printing behavior equipment. Additionally, we thank the Versiti Blood Research Institute for providing access to their Shared Imaging Facility. Finally, we would like to thank Aniko Szabo, PhD, of the Medical College of Wisconsin Biostatistics Department for providing data analysis advice.

## Author Contributions

A.R.M. contributed to the study conception and design, planned, performed, and analyzed behavioral and calcium imaging experiments, and wrote and edited the manuscript. E.I. contributed to the study conception and design, planned, performed, and analyzed calcium imaging and electrophysiological experiments, designed figures, and wrote and edited the manuscript. M.L.S. planned and performed calcium imaging experiments and edited the manuscript. A.D.M. planned, performed, and analyzed immunohistochemistry experiments and edited the manuscript. A.S. planned, performed, and analyzed qRT-PCR experiments, edited the manuscript. M.M.P. contributed to the design and optimization of calcium imaging experiments, performed, and analyzed calcium imaging experiments, and edited the manuscript. S.M.S. contributed to the design of the HEK cell experiments, assisted with optimizing transfection protocols, and edited the manuscript. K.E.S. contributed to the study conception and design and edited the manuscript. H.Y. contributed to the design of the HEK cell experiments, assisted with optimizing transfection protocols, and edited the manuscript. C.L.S. contributed to the study conception and design and in editing the manuscript.

## SUPPLEMENTAL MATERIALS

**Supplementary Fig. 1.**
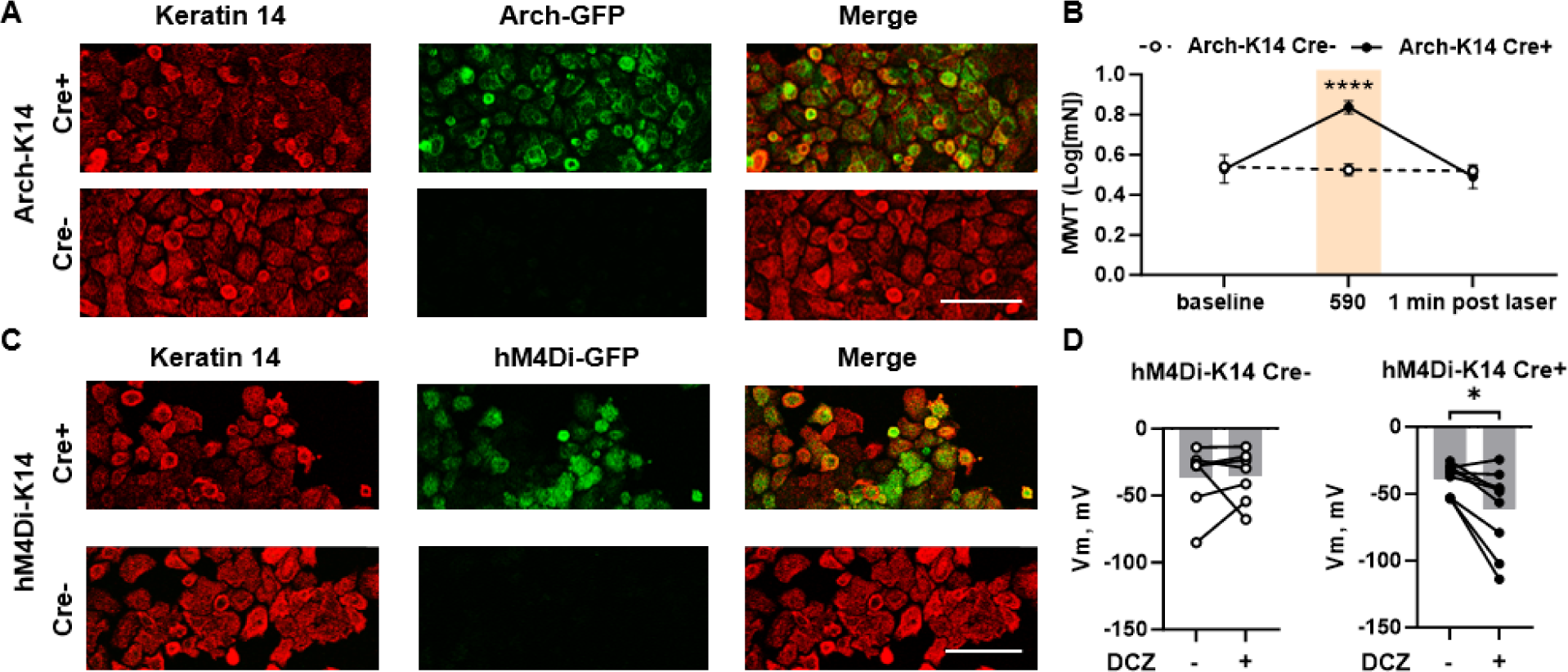
Validation of optogenetic and chemogenetic mouse lines. (**A**) Cultured mouse keratinocytes from Arch-K14 Cre- and Arch-K14 Cre+ animals. Immunoreactivity to keratin 14 is shown in red, while immunoreactivity for GFP is shown in green. Overlay data shows that the Arch-K14 Cre-animal has no GFP expression in K14-expressing keratinocytes. (**B**) Withdrawal thresholds of naïve Arch mice return to baseline levels 1 min after 590 light is removed; n = 4-5 animals. (**C**) Cultured mouse keratinocytes from hM4Di-K14 Cre- and hM4Di-K14 Cre+ animals. Immunoreactivity to keratin 14 is shown in red, while immunoreactivity for GFP is shown in green. Overlay data shows that the Arch-K14 Cre-animal has no GFP expression in K14-expressing keratinocytes. (**D**) Changes of hM4Di-K14 Cre- and hM4Di-K14 Cre+ keratinocytes’ membrane potential upon 10 min treatment with deschloroclozapine (DCZ, 0.5 µM). Dots represent individual cells; n = 3 animals. Statistical analysis for (**C**) was performed using 2way ANOVA with Tukey’s multiple comparisons tests; for (**D**) using paired Student’s *t* test. Error bars, ± SEM. *P < 0.05, **P < 0.01.

**Supplementary Fig. 2.**
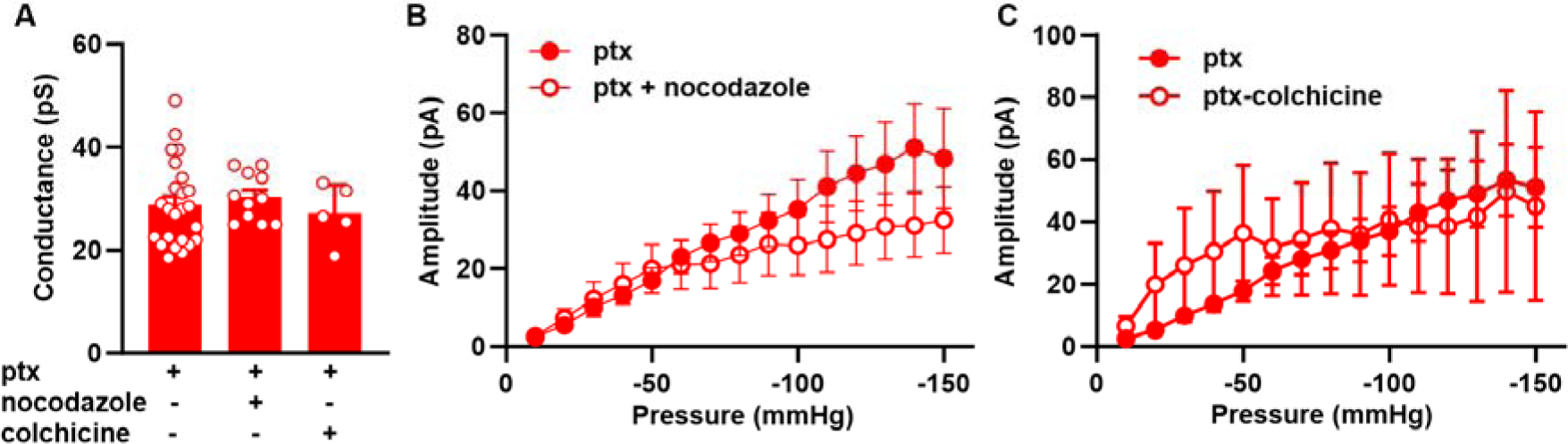
Nocodazole and colchicine do not reverse the paclitaxel-induced sensitization of Piezo1 currents. **(A)** Summary data of the effect of pretreatment of Piezo1 expressing HEK-P1KO cells with paclitaxel, paclitaxel + nocodazole, and paclitaxel + colchicine on Piezo1 conductance. Current-pressure relationship of stretch-activated currents recorded at a V_hold_ = -80 mV from paclitaxel (n=39), paclitaxel + nocodazole (**B**, n=21), and paclitaxel + colchicine (**C**, n=6) treated HEK-P1KO cells expressing Piezo1. Error bars, ± SEM.

